# Novel cancer subtyping method based on patient-specific gene regulatory network

**DOI:** 10.1101/2021.03.24.436731

**Authors:** Mai Adachi Nakazawa, Yoshinori Tamada, Yoshihisa Tanaka, Marie Ikeguchi, Kako Higashihara, Yasushi Okuno

**Author notes:** Innovation Center for Health Promotion, Hirosaki University (Yoshinori Tamada).

## Abstract

The identification of cancer subtypes is important for the understanding of tumor heterogeneity. In recent years, numerous computational methods have been proposed for this problem based on the multi-omics data of patients. It is widely accepted that different cancer subtypes are induced by different molecular regulatory networks. However, only a few incorporate the differences between their molecular systems into the classification processes. In this study, we present a novel method to classify cancer subtypes based on patient-specific molecular systems. Our method quantifies patient-specific gene networks, which are estimated from their transcriptome data. By clustering their quantified networks, our method allows for cancer subtyping, taking into consideration the differences in the molecular systems of patients. Comprehensive analyses of The Cancer Genome Atlas (TCGA) datasets applied to our method confirmed that they were able to identify more clinically meaningful cancer subtypes than the existing subtypes and found that the identified subtypes comprised different molecular features. Our findings show that the proposed method, based on a simple classification using the patient-specific molecular systems, can identify cancer subtypes even with single omics data, which cannot otherwise be captured by existing methods using multi-omics data.

## Introduction

Cancer is a highly heterogeneous disease and is known to differ among patients. This heterogeneity renders one cancer type to be composed of multiple subtypes, which are characterized by different molecular features. Clinical identification of these cancer subtypes is currently one of the major challenges in cancer research. Identifying the subtypes can provide an understanding of the underlying molecular mechanisms and thereby design precise treatment strategies for efficient cancer management. In recent years, advances in high-throughput sequencing technologies have generated large amounts of data on various cancer types. For example, The Cancer Genome Atlas (TCGA) contains multi-omics data, including gene expression, mutation, methylation, and copy number, of over 34 cancer types. These multi-omics data allow improvements in cancer subtyping via computational methods^1–3^. However, most studies do not classify the cancer subtypes based on differences in the molecular systems, but they are based only on the differences in the numerical patterns of the omics data.

Network representation of molecule-to-molecule relationships is a key to understanding a fundamental molecular system, and it plays an important role in understanding each biological process and the molecular mechanisms of cancer^4^. Therefore, knowledge of such networks could be a promising data source for cancer subtyping. Some well-known types of biological networks are gene regulatory networks and protein–protein interaction networks^5^. Although the importance of molecular systems has been shown in recent years, only a few studies have incorporated the knowledge of molecular networks into their clustering processes^6–8^. However, these methods do not sufficiently express the molecular systems for two reasons. First, the networks used do not contain a large number of genes that are supposed to be expressed in cells. In fact, only those genes that are already known to be involved in certain cancer types have been included in the networks^6–8^. Second, the networks used are constructed from public databases that do not include condition-dependent networks^9,10^. Recent studies have revealed that biological networks vary between normal and the disease states^11,12^. Because genetic interactions are condition-specific, the networks of particular types of cancers are different from those found in these databases.

In this study, we propose a novel method to classify cancer subtypes by incorporating differences in the molecular systems of the patients. Because a gene network involves gene–gene regulatory relationships and is a fundamental network among the various molecular networks, it can be an adequate representation of molecular systems for our purpose. Therefore, our proposed method is based on the estimated gene network from the gene expression data of patients. The main scheme of the proposed method is illustrated in Fig. 1. Briefly, our method estimates a gene network from a gene expression dataset in the tumors of patients using a Bayesian network. A numerical value is then calculated for every edge of the estimated gene network with respect to each patient’s sample. This edge value, also known as the edge contribution value (ECv), is derived by evaluating the contribution of the edge to an expression value with respect to a patient in terms of the estimated molecular system^13^. Therefore, differences in ECvs reflect differences in the molecular systems of particular patients. This calculation of ECvs generates a matrix of numerical values consisting of patient-specific networks. Finally, hierarchical clustering was performed using the matrix to classify patients into subtypes. This simple clustering allows the identification of various subtypes, incorporating complex patient-specific activities of their molecular systems, which cannot be captured by the existing classification methods using the multi-omics data of patients. We used this method to analyze two cancer types from TCGA datasets, namely stomach adenocarcinoma and lung cancer, including lung adenocarcinoma and lung squamous cell carcinoma. Consequently, all two cancer types were classified into three major novel subtypes, which defined their differential prognoses and distinct molecular properties. Our method identified system-based cancer subtypes using only transcriptome data, which is more accessible compared to other omics data. Additionally, the proposed method allowed for the extraction of subnetworks to explain the features of the identified subtypes. Collectively, our findings indicate that the proposed method can successfully incorporate cancer-specific gene networks and establish a novel cancer subtype classification that overcomes the limitations of other sophisticated clustering methods, which are based on gene expression data alone.

**Figure 1.**
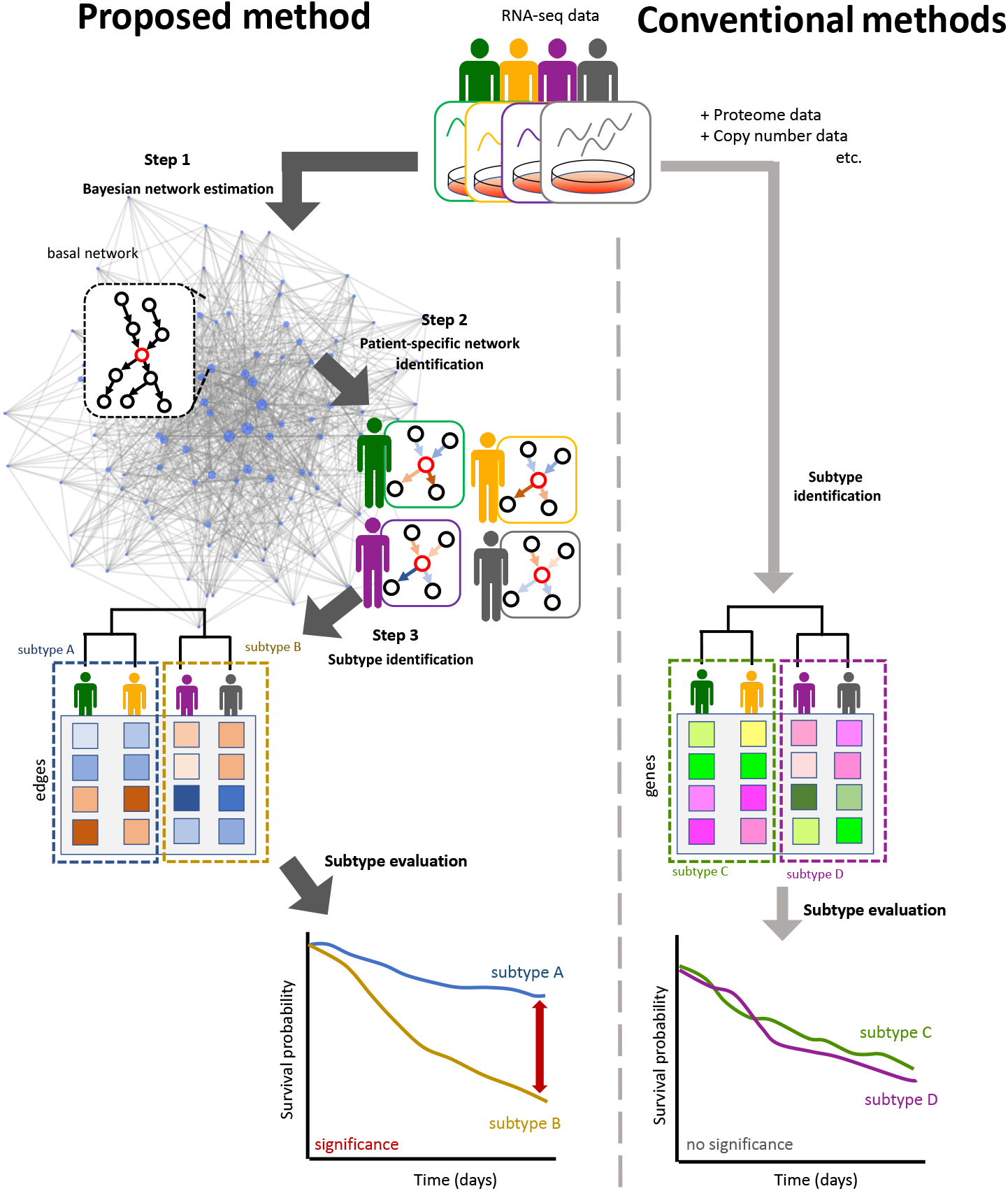
Overview of our method.

## Methods

In this chapter, we first introduce the gene network estimation method using a Bayesian network. We then explain the ECvs that allow us to quantify the patient-specific characteristics of the gene networks. Finally, the method for subtype classification of cancer patients based on their ECvs is described.

### Bayesian network with *B*-spline nonparametric regression

To define the transcriptomic molecular networks, or gene networks, a Bayesian network with *B*-spline nonparametric regression model is used^14^. A Bayesian network (BN) is a graphical model that represents the cause-and-effect relations among variables as a directed acyclic graph. By representing the gene expressions as the random variables, we can estimate the system-level regulatory relationships from the transcriptome data. Many successful studies have reported on the use of BN for gene network analysis^15–18^. Assuming *p* genes, the joint density of the gene expressions in a BN is described as

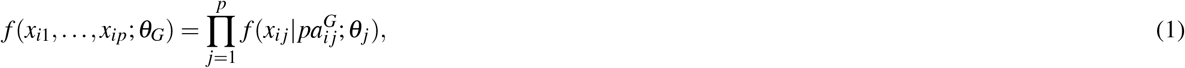

where *x_ij_* represents the gene expression of the *j*-th gene at the *i*-th sample, *θ_G_* is the parameter vector of the BN represented by 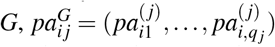 denotes the gene expression vector of *q_j_* parents of the *j*-th gene, and *θ_j_* is the parameter vector for the local density with respect to the *j*-th gene. The optimal structure of the network is obtained by the maximization of the posterior probability given the observed data as

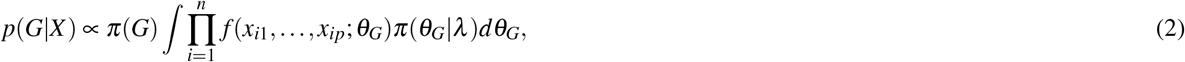

where *X* is the observed data matrix, *π*(*G*) is the prior probability of *G, n* is the number of samples in *X*, *π*(*θ_G_*|*λ*) denotes the prior distribution of *θ_G_*, and *λ* is the hyperparameter vector. The drawback of the BN is that obtaining the optimal structure for a given dataset is NP-hard. Therefore, the neighbor node sampling and repeat (NNSR) algorithm was used^19^.

### Classification of patients based on their molecular networks

Tanaka et al. (2020)^13^ proposed the edge contribution value (ECv) to extract subnetworks from Bayesian networks, related to specific differences observed for in vitro experiments. Briefly, the *B*-spline nonparametric BN assumes that the gene expression is modeled as

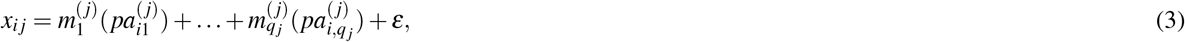

where 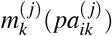 is a regression function using *B*-spline curves for the *k*-th parent of the *j*-th gene, and *ε* is the error term. Because a value of this regression function 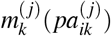 can be considered as a contribution of an edge from the *k*-th parent to the *j*-th gene, Tanaka et al. (2020)^13^ defined ECv as

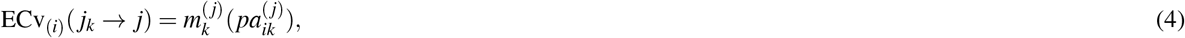

where *j_k_* represents the index of the *k*-th parent of the *j*-th gene. Tanaka et al. (2020)^13^ considered the differences of ECvs as ΔECv between the control and the TGF*β*-treated samples, which extract the distinctive edges with a certain threshold for ΔECv. These were defined as the subnetworks characterizing the EMT in lung cancer cell lines. Here, we propose an algorithm that uses ECvs to characterize the patients and elucidate cancer subtypes. Using ECvs as quantified gene networks, patients with similar molecular systems would have similar ECvs, while patients with different molecular systems would have different quantified networks. In this context, clustering based on the molecular system differences enables us to identify cancer subtypes. This requires the gene network estimation from the gene expression data of patients and the calculation of ECvs values for the estimated edges.

### Classification of cancer subtypes

Assuming *E* edges in the estimated gene network, the patient’s quantified network is defined as a vector of *E* elements (*c*_*i* 1_,…,*c_iE_*), where *c_iv_* is an ECv of the *v*-th edge for the *i*-th patient. The quantified networks of all the patients are collected and used to construct the columns of a matrix, resulting in an ECv matrix whose (*i, v*) element corresponds to an ECv of the *v*-th edge for the *i*-th patient. Using clustering, the patients of this ECv matrix are classified according to the differences and similarities between their gene networks. The ECv matrix consists of all the edges of the network, including approximately 20,000 genes and 150,000 edges. Since the majority of the edges do not represent differences in terms of ECv, parts of the edges are selected prior to clustering. To select the edges for hierarchical clustering based on the ECv matrix, the variance of each edge among patients is used as the ranking edges to represent the differences across the samples. The top *N* edges showing large variances will be selected. Therefore, hierarchical clustering is performed for the ECv matrix, consisting of the selected *N* edges, for the classification of patients into the different cancer subtypes.

### Extraction of edges

Although hierarchical clustering classifies patients into cancer subtypes, the part of the network that is affected by the clustering result is unknown. Therefore, distinctive edges with significant ECv differences need to be extracted, as in Tanaka et al. (2020)^13^. In their study, they extracted the edges by calculating the ΔECv between two conditions. However, since their method cannot be applied for more than two groups, we extended their scheme. The following proposed method allows us to extract distinctive edges with significant ECv differences using ECvs in multiple groups. Suppose that there are *M* groups of patients *R*_1_,…, *R_M_*. We define 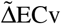 with respect to group *R_r_* out of these *M* groups as

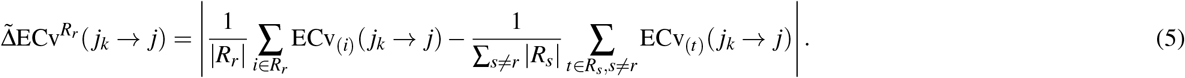

The 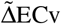 of every single edge is then calculated with respect to each subtype, where significant 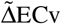 edges are regarded as distinctive edges of specific subtypes.

## Results

### Dataset

In this study, our proposed method was applied to TCGA RNA-seq datasets of three types of cancer: stomach adenocarcinoma (STAD)^20^; lung cancer, including lung adenocarcinoma (LUAD)^21^ and lung squamous cell carcinoma (LUSC)^21^. LUAD and LUSC were regarded as one dataset and referred to LUNG as they are both lung cancers, and we test whether they are split into different subtypes. These datasets were preprocessed as described in Supplementary information (Supplementary S2.1).

### Classification based on ECv matrix

We calculated the ECv of every single edge in the estimated network for each patient, respectively. This ECv calculation generated a matrix of numerical values consisting of patient-specific molecular systems. To select the edges for hierarchical clustering based on the ECv matrix, the variances of edges were calculated as described in Method section. For TCGA datasets, we selected the top *N* = 250 edges with the highest variances in ECv among the patients. These 250 edges corresponded to approximately 0.01% of the total number of edges. Classifying the patients by the small number of edges supposed to be a potentially better classification method. To classify the patients into network-based subtypes, we performed hierarchical clustering for the ECv matrix consisting of the selected edges in three types of cancer. The ECv heatmap revealed a high variance among the intrinsic ECv matrix of the samples (Fig. 2a, Fig.S1). The clustering results of the ECv heatmap indicate that patients of each cancer type were classified into three major subtypes, namely subtype 1, subtype 2, and subtype 3, according to the similarities and differences between the patient networks (Fig. 2a, Fig.S1). In our dataset, *N* = 250 was approximately the minimum number of edges that produced biologically and clinically meaningful results, as we described in Result section later.

**Figure 2.**
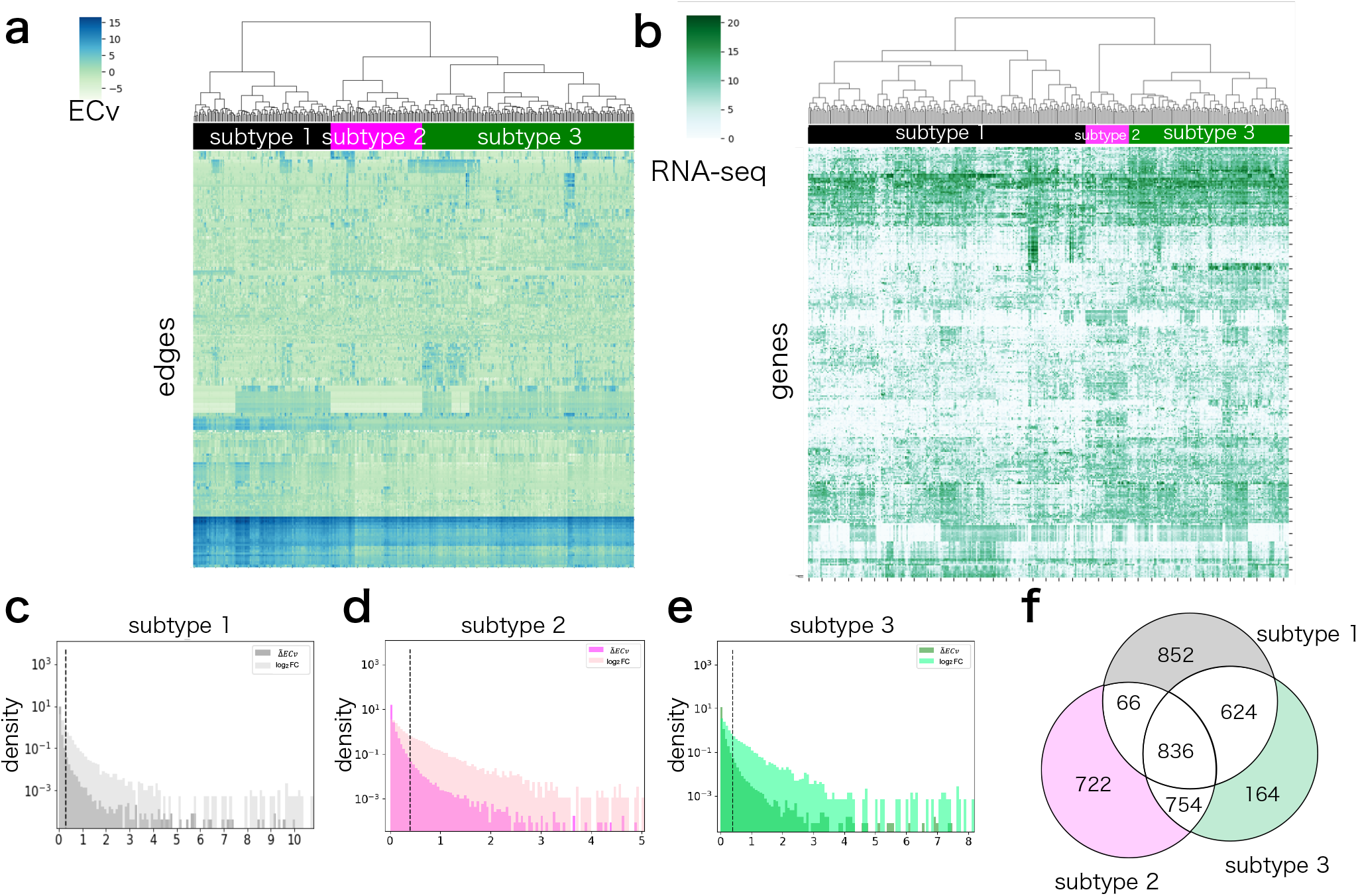
(a) Heatmap showing hierarchical clustering for the ECv matrix in the STAD dataset. (b) Heatmap showing hierarchical clustering for the RNA-seq matrix in the STAD dataset.(c-e) The distribution of ΔECv of edges and absolute log_2_ fold change in genes in the STAD dataset (See supplementary S2.3). Dashed lines represent of the top 1.0% of the total edges in every subtype. (f) The Venn diagram represents the number of edges in the STAD dataset. Colored areas in the Venn diagram represent subtype-specific edges in each subtype.

### Extraction subtype-specific edges

The ΔECv value was calculated for every single edge in each subtype across three types of cancer. Edges with a high represent significant differences between subtypes. The distribution of suggested that only limited edges showed significant differences (Fig. 2c-e, Fig. S2). Based on the distributions of 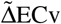, the top 1.0% of the total edges in the estimated network were found to differ significantly. Therefore, we selected the corresponding edges from each subtype and removed any edges that were also selected for other subtypes (Fig. 2f, Fig. S3). We denoted the extracted edges as subtype-specific edges. Networks consisting of these subtype-specific edges were considered as a subnetwork characterizing the identified subtypes.

### Applications in TCGA stomach cancer datasets

Stomach cancer is one of the most common leading causes of cancer-related death worldwide^22^. In the original TCGA paper, the authors demonstrated that stomach cancer is a heterogeneous disease with four molecular subtypes—Epstein-Barr virus (EBV), microsatellite instability (MSI), genomically stable (GS), and chromosomal instability (CIN)^20^. These subtypes are based on the six platforms of the multi-omics molecular signature: somatic mutation, mRNA expression, miRNA expression, promoter methylation, somatic copy number alteration, and protein expression^20^. However, as previously mentioned^20^, no significance was observed between the prognoses of the subtypes (log-rank test *p*-value = 0.10 > 0.05) (Fig. 3a). This suggests that the multi-omics-based subtypes did not account for the clinical significance, such that subtyping may not provide an opportunity to improve therapeutic treatments. We hypothesized that multi-omics data provide limited information on tumor subtyping. Rather, the differences in molecular systems may explain the differences in patients’ prognoses.

**Figure 3.**
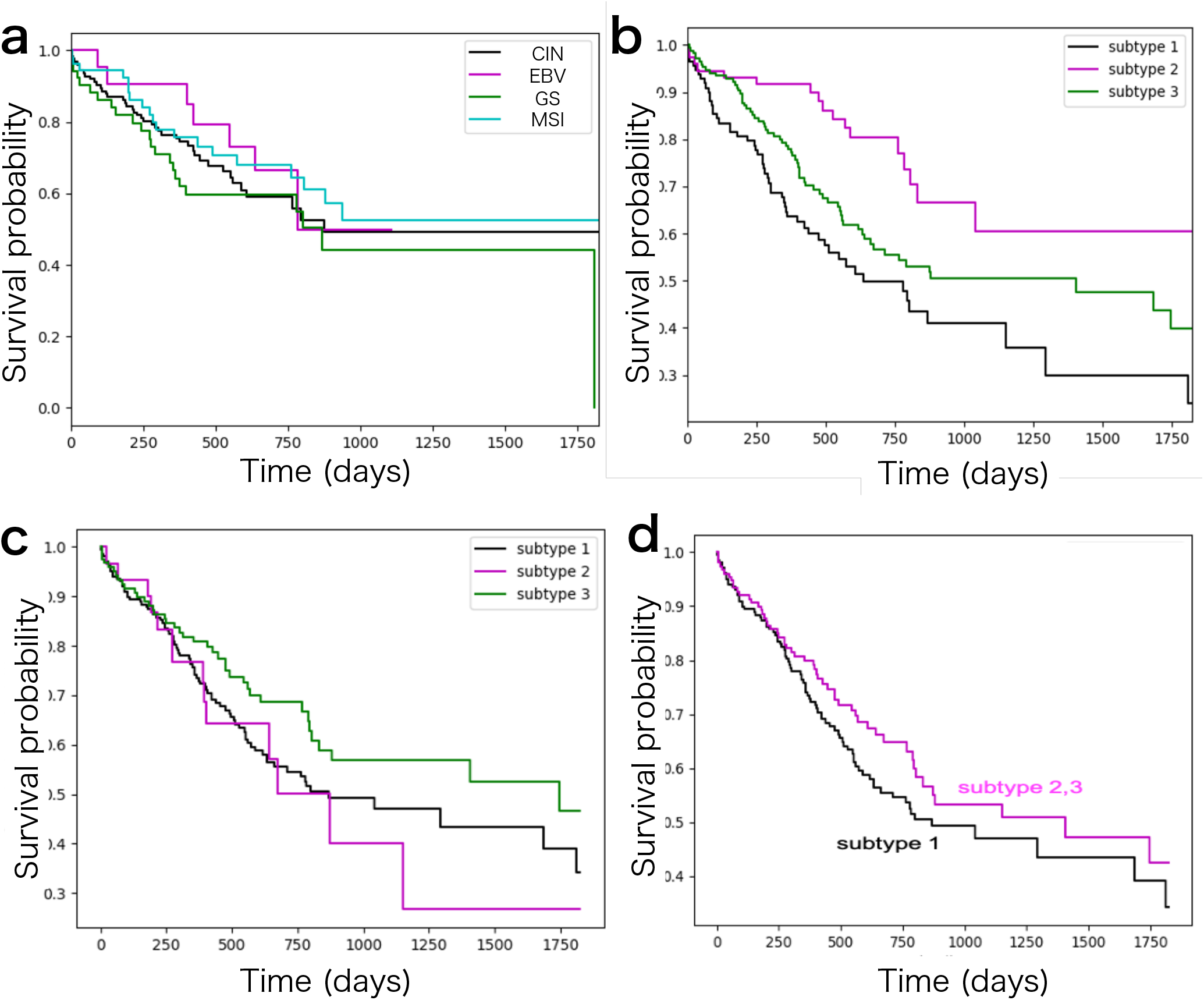
(a) Kaplan-Meier survival probability curves of patients for the multi-omics-based subtypes. The log-rank test between two subtypes; 0.45 (CIN vs EBV) > 0.05, 0.30 (CIN vs GS) > 0.05, and 0.50 (CIN vs MSI) > 0.05, 0.31 (EBV vs GS) > 0.05, 0.95 (EBV vs MSI) > 0.05, 0.16 (GS vs MSI) > 0.05. (b) Kaplan-Meier survival probability curves of patients for the identified network-based subtypes. The log-rank test between two subtypes; 0.00016 (subtype 1 vs 2) < 0.05, 0.042 (subtype 1 vs 3) < 0.05, and 0.013 (subtype 2 vs 3) < 0.05. (c) Kaplan-Meier survival probability curves of patients for the identified RNA-seq based three subtypes. The log-rank test between two subtypes; 0.70 (subtype 1 vs 2) > 0.05, 0.091 (subtype 1 vs 3) > 0.05, and 0.14 (subtype 2 vs 3) > 0.05. (d) Kaplan-Meier survival probability curves of patients for the identified RNA-seq based two major subtypes. The log-rank test *p*-value = 0.19 > 0.05

To address this issue, we applied the proposed method to the preprocessed RNA-seq datasets of STAD. As described in Result section above, the clustering results of the ECv heatmap indicate that stomach cancer is classified into three major subtypes: subtype 1 (113 samples), subtype 2 (76 samples), and subtype 3 (173 samples) (Fig. 2a). To determine the relationship between the existing multi-omics-based subtypes and our identified subtypes, we summarized the number of patients across them (Table 1) and found that our subtyping was different from the multi-omics-based subtypes. These findings suggest that our proposed method, based on the patient-specific molecular systems, can identify novel cancer subtypes that cannot be captured by existing methods using multi-omics data. To investigate the extent to which our proposed method classifies cancer subtypes, we conducted a survival analysis of the three identified subtypes. A better method for subtype classification is key for the identification of cancer subtypes and different prognoses, since patients with different molecular systems require different drug treatments. The Kaplan-Meier survival probability curves in the identified subtypes indicated that each subtype had a significantly different prognosis pattern (log-rank test *p*-value = 0.00011 < 0.05) (Fig. 3b).

**Table 1.** The relationship between the existing four molecular subtypes and our identified subtypes.

| subtype name | CIN | EBV | GS | MSI | Unknown | All |
| --- | --- | --- | --- | --- | --- | --- |
| subtype 1 | 33 | 10 | 37 | 9 | 24 | 113 |
| subtype 2 | 25 | 3 | 5 | 21 | 22 | 76 |
| subtype 3 | 58 | 11 | 6 | 15 | 83 | 173 |

Furthermore, to confirm whether the gene network information improves the classification of the cancer subtypes, hierarchical clustering was performed using RNA-seq data alone, without network information. The top 322 genes showing the highest variances of the RNA-seq data in STAD were selected, as the 250 edges with the ECv matrix were composed of 322 genes. Consequently, we identified three RNA-seq-based subtypes (Fig. 2b). However, these subtypes did not show any significant differences in terms of their prognoses (Fig. 3c). Despite employing two major subtypes in the clustering result, the differences were not significant (Fig. 3d). To determine the relationship between the network-based and the RNA-seq-based subtypes, we summarized the number of patients across them (Table S2) and found that network-based subtypes were different from the RNA-seq-based subtypes. These results further suggest that our network-based method might generate a better cancer subtyping profile. Moreover to confirm whether the gene network information improves the classification of the cancer subtypes, we also applied the iNMF method, since it is a successful method for cancer subtyping that can handle multi-omics data^23^. We set three clusters when performing the iNMF as we identified three subtypes in our method. However, although we performed using gene expression data alone and using multi-omics data consisting of gene expression, miRNA expression, copy number and DNA methylation, in both cases, these subtypes did not show any significant differences in their prognosis (Fig.S4).

To characterize the network-based subtypes obtained from the ECv matrix, we highlighted the subnetworks composed of subtype-specific edges in the estimated basal network (Fig. 4a). The subtype-specific edges constituted a module in the basal network, especially in terms of subtype 3 (Fig. 2f, Fig. 4a). In Fig. 4a, the node layout of the basal network was arranged only using its topological structure. This suggests that the differences in the partial modules of the network might affect the classification of the cancer subtypes. Furthermore, to account for the properties of the identified subtypes, we verified their molecular features using gene ontology analysis. The subtype-specific networks were composed of 250, 1186, and 407 genes in subtype 1, subtype 2, and subtype 3, respectively. The ontology analysis results indicated that, according to the top five biological function terms, each subtype had a characteristic molecular feature (Table 2). Although most of the biological functions in the subtypes were related to development, the developmental stages or tissues varied between the subtypes. For example, “cardiovascular system development and function” was found in subtype 1, while “embryonic development” was found in subtype 2 and “cellular development” was found in subtype 3. In particular, the top five of biological functions in subtype 3, which were associated with a moderate prognosis, were completely different from those in the other subtypes. Most of the biological functions observed in subtype 1 and subtype 2 were related to development, while “cellular growth and proliferation” and “cell-to-cell signaling and interaction” were observed exclusively for subtype 3. Furthermore, we visualized a network composed of subtype-specific edges (Fig. 2f) and extracted the largest connected component from the visualized network in each subtype (Fig. 4b). The results suggested that the subtype-specific edges in subtypes 2 and 3 were composed of a large connected subnetwork. Moreover, we found *SALL2, ETNK2*, and *APBB1* were located as the top hub genes in subtype 2. These genes are implied as cancer-related genes^24–26^, and thus may play an important role in characterizing these subtypes.

**Figure 4.**
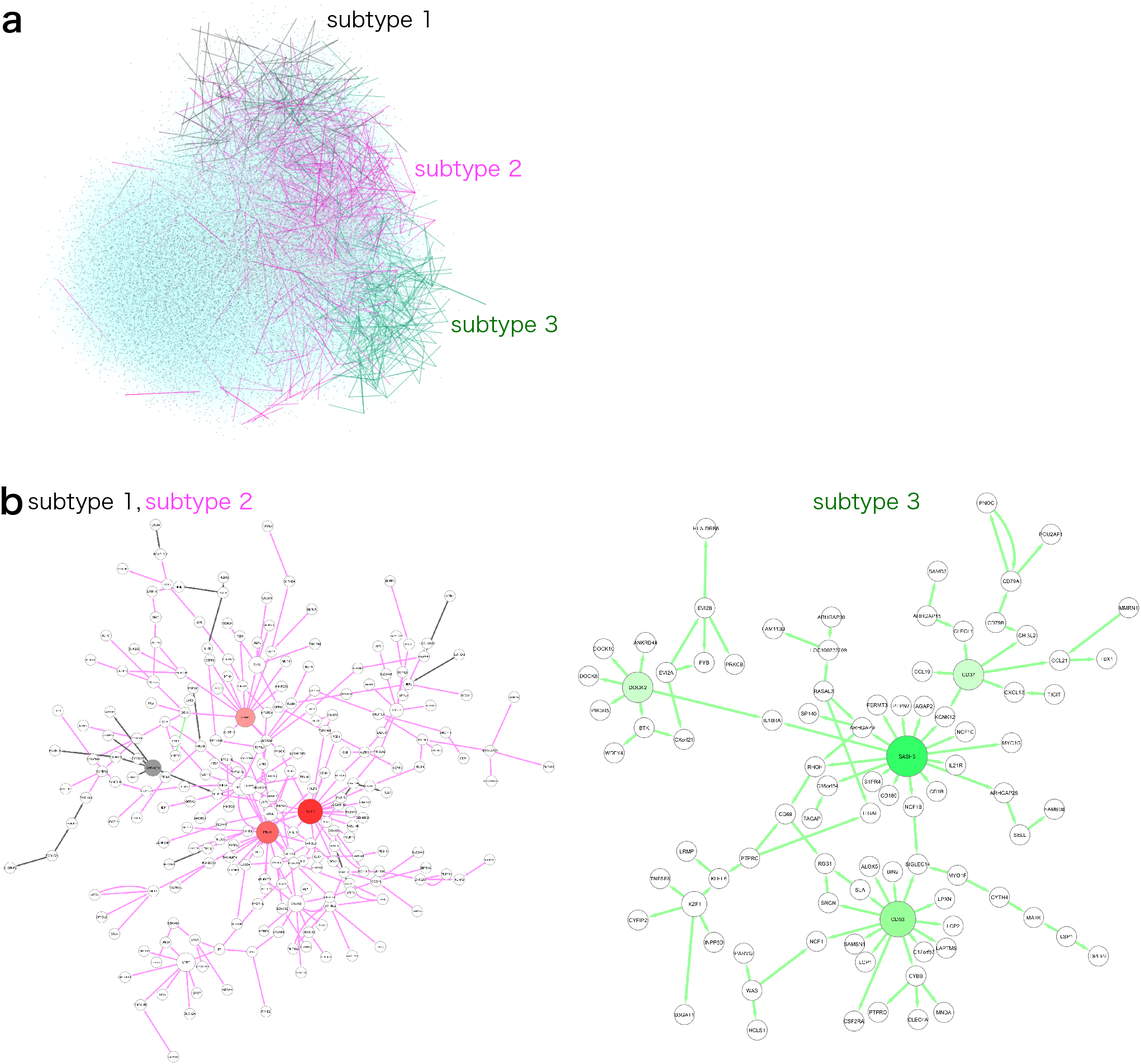
Visualization of subtype-specific subnetworks in the STAD dataset. (a) Subnetworks of subtype-specific edges were highlighted with the basal network (blue). (b) The biggest connected component in the subnetwork of subtype-specific edges in each subtype. Edges and nodes were colored by each subtype: subtype 1 (gray), subtype 2 (magenta), and subtype 3 (green). Colored nodes were hub nodes in each subtype and the color gradient represents the outdegree of hubs.

**Table 2.**
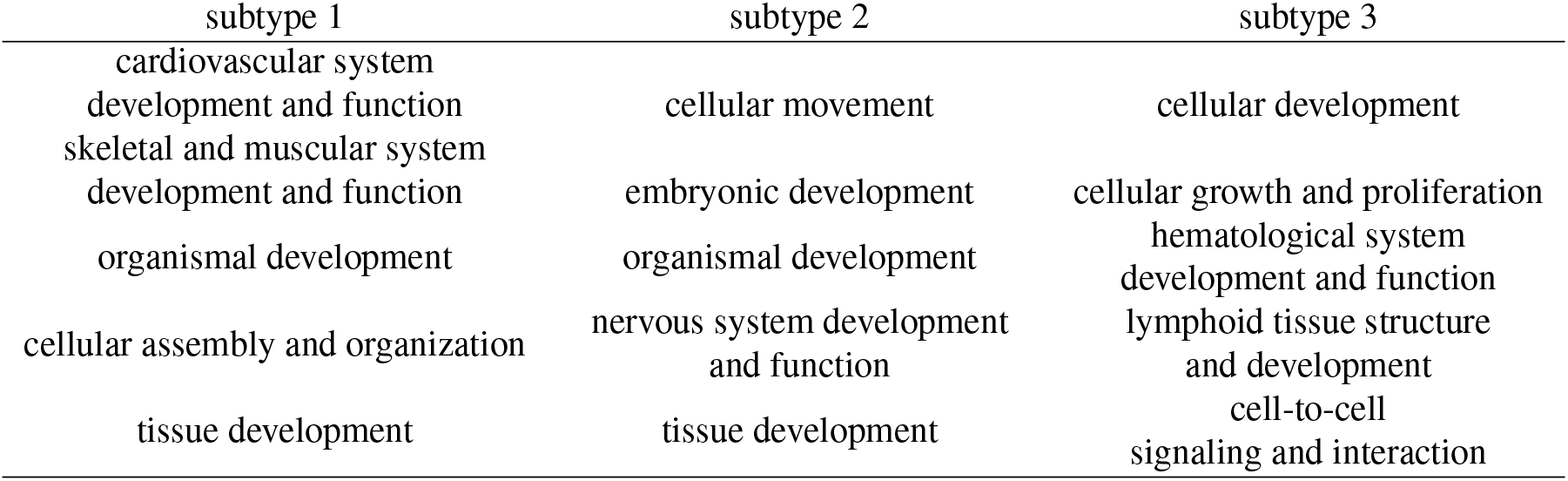
The top five terms of biological functions in the STAD dataset.

### Identification of cancer subtypes in lung cancer

To test the effectiveness of our method in other datasets, we applied it to the lung cancer dataset (LUNG). As shown in Fig. S1, the clustering results of the ECv heatmap indicate that lung cancer is classified into three subtypes: subtype 1 (227 samples), subtype 2 (121 samples), and subtype 3 (343 samples). Then, survival analysis was conducted, similar to STAD analysis. The Kaplan-Meier survival probability curves in the identified subtypes indicate that each subtype has a significantly different pattern of prognosis (log-rank test *p*-value = 1.3e-14 < 0.05) (Fig. S5a). Next, the differences in the molecular features between the identified subtypes were determined. The subtype-specific networks were found to be composed of 158, 1049, and 582 genes in subtype 1, subtype 2, and subtype 3, respectively. Gene ontology analysis of the subtypes indicate that all of the top five biological functions varied between them (Table S3). These findings suggest that our proposed method might also work for different cancer types. Moreover, hierarchical clustering followed by survival analysis, was performed for RNA-seq data as shown in the STAD section. Consequently, three RNA-seq-based subtypes were identified and the prognoses of these subtypes were significantly different (log-rank test *p*-value = 1.5e-15 < 0.05) (Fig. S5b and S5c). While the patients in network-based subtype 1 were almost identical with those in RNA-seq-based subtype 1, those in network-based subtype 2 and subtype 3 were different from RNA-seq-based subtypes (Table S4). Furthermore, the network-based and RNA-seq-based clustering could almost completely classify LUAD and LUSC (Table S4). This may suggest that patients with various molecular features can be classified even without network information, as LUAD and LUSC have characteristic molecular features that are significant for classifying them using transcriptome data^27, 28^. Gene ontology analysis indicates that network-based subtypes could reveal completely different characteristic molecular features among the subtypes. Thus, this suggests that our method can identify novel subtypes that cannot be detected using RNA-seq clustering. We also visualized a network composed of subtype-specific edges (Fig. S6a) and extracted the largest connected component from the visualized network in each subtype (Fig. S6b-d).

## Discussion

In this study, we proposed a novel method for the classification of cancer subtypes based on patient-specific molecular systems. The proposed method is able to identify novel subtypes with different prognoses, as well as the differences in molecular properties between stomach cancer and lung cancer. Differences in molecular systems are not necessarily associated with the prognoses of patients. However, it is likely to affect the effectiveness and/or medical treatment options available for these patients. For this reason, our novel subtypes may be related to prognosis of patient.

Although many types of omics data are currently available, it remains difficult to integrate multi-omics data in research. Each type of omics data can be used to classify cancers into various subtypes in terms of prognosis, pathological findings, and others. However, since our proposed method uses only transcriptome data, even though our gene network-based method was successful, it may not be sufficient to obtain an in-depth understanding of the molecular systems. Despite this, changes in the different layers of omics networks influence the transcriptome profile at some level. This could explain why our proposed method, based on the gene network, was able to identify novel cancer subtypes using only the transcriptome data. There, however, remains room for improvement in the method reported in this study, wherein classification using multi-omics data based on estimated systems represents an informative strategy for the identification of cancer subtypes.

## Data availability

All the patient lists generated in this study are provided in the supplementary data. All the networks are available at NDEx (The basal network in STAD; https://www.ndexbio.org/viewer/networks/1dabd135-8bab-11eb-9e72-0ac135e8bacf, The subtype-specific network in STAD; https://www.ndexbio.org/viewer/networks/4e61c7cf-8889-11eb-9e72-0ac135e8bacf, The basal network in LUNG; https://www.ndexbio.org/viewer/networks/0e943431-8e00-11eb-9e72-0ac135e8bacf, The subtype-specific network in LUNG; https://www.ndexbio.org/viewer/networks/be019ba4-8e01-11eb-9e72-0ac135e8bacf).

## Acknowledgements

We thank Dr. K. Fukuyama for advice on the clinical interpretation and members of the Okuno Lab. and Project 20 in Life Intelligence Consortium for their helpful discussions. Computational resources were provided by the Super Computer System, Human Genome Center, Institute of Medical Science, University of Tokyo. This work was supported by Cabinet Office, Government of Japan, Public/Private R&D Investment Strategic Expansion Program (PRISM). This work was supported by RIKEN Junior Research Associate Program.

## Author contributions statement

M.A.N, Y.T. (Yoshinori Tamada),Y.O. conceived the experiments, analyzed the result, and wrote the manuscript; Y.T. (Yoshihisa Tanaka), M.I., and K.H. conducted experiments; Y.T. (Yoshihisa Tanaka) and Y.O. reviewed and edited the manuscript.

## Compete interests

Y. Tamada and Y. Okuno have a patent application on the method for identification of patient-specific network used in this study through the technology licensing organization in Kyoto University. Other authors declare no conflict of interest.

