## Supplemental document for "Novel cancer subtyping method based on patient-specific gene regulatory network"

### Novel cancer subtyping method based on patient-specific molecular networks

All supplementary data is available at https://ytlab.jp/suppl/nakazawa2021/index.html

#### Supplement Information

#### S1 Methods

##### S1.1 Computational and statistical environment

All the network estimations and ECv calculations were performed using the SHIROKANE supercomputer system at Human Genome Center, the Institute of Medical Science, the University of Tokyo.

All statistical analyses were performed using Python unless stated otherwise.

Hierarchical clustering was performed using the "ward" methods and Euclidean distance in Python library. The survival analysis was evaluated using the log-rank test in R package survival and Python library lifelines. Molecular function analysis was performed using Ingenuity Pathway Analysis (IPA)[1]. Network visualization and analysis were performed in Cytoscape[2].

#### S2 Result

##### S2.1 Dataset

These RNA-seq data were downloaded from UCSC Xena[3]. Clinical data were downloaded from GDC Data Portal at TCGA to evaluate the results of the analysis. Patients were selected for whom both the RNA-seq data of the tumor specimens and the clinical data (five years) for each cancer type were available. Next, the genes with a mean percentile under 15 were removed from the RNA-seq data of each dataset. Ultimately, 365 patients were selected for STAD and 692 patients for LUNG. The preprocessed RNA-seq datasets comprised different sets of 17,450 genes.

##### S2.2 Network estimation

The NNSR algorithm determines the final network structure by extracting the edges whose estimated frequencies are greater than a given threshold. A threshold of 0.1 was used in our analysis, as in Tanaka et al. (2020)[4]. This algorithm repeatedly estimates subnetworks, including a thousand nodes, such that we first checked whether the algorithm produced stable networks. The network estimation was conducted independently three times, and the concordance of the estimated edges was calculated between the two estimated networks as an indicator of robustness in the estimated networks, as in Tanaka et al. (2020)[4]. Consequently, when the number of iterations ($T$) of the subnetwork estimation was 100,000, as recommended by Tamada et al. (2011)[5], the concordance was less than 95%. However, since this is slightly below the necessary level of network stability, we used $T = 300,000$ and obtained a concordance of 96.3% and 95.6%, for STAD and LUNG, respectively (Table S1). These results suggested that the structures of the estimated networks were stable, and that $T$=300,000 was sufficient for our analysis.

##### S2.3 Comparison of the $\tilde{\Delta}ECv$ and FC distribution

The distributions between $\tilde{\Delta}ECv$ and $\log_2$ fold change (FC) were compared to present that only the limited edges show significant differences. The distribution of $\tilde{\Delta}ECv$ is much steeper than that of $\log_2$ FC. The FC for RNA-seq data is defined as one subtype/the rest of two subtypes. We overlapped $\tilde{\Delta}ECv$ and FC in the same histogram (Fig. 2c-e, Fig. S2).

##### S2.4 Hierarchical clustering for RNA-seq data

For LUNG dataset, the top 310 genes showing the highest variances of the RNA-seq data were selected for hierarchical clustering, as the 250 edges with the ECv matrix in LUNG were composed of 310 genes.

**Table S1.** Concordance of the estimated networks.

| | $T = 100,000$ | $T = 300,000$ |
|---|---|---|
| STAD | 92.7% | 96.3% |
| LUNG | 92.7% | 95.6% |

**Table S2.** The summarization of the number of patients across subtypes identified by the clustering of ECv matrix and RNA-seq data.

| | | RNA-seq | | |
|---|---|---|---|---|
| | | subtype 1 | subtype 2 | subtype 3 |
| ECv | subtype 1 | 91 | 8 | 15 |
| | subtype 2 | 43 | 0 | 33 |
| | subtype 3 | 77 | 24 | 74 |

**Table S3.** The top five terms of biological functions.

| cancer type | subtype 1 | subtype 2 | subtype 3 |
|---|---|---|---|
| STAD | cardiovascular system development and function | cellular movement | cellular development |
| | skeletal and muscular system development and function | embryonic development | cellular growth and proliferation |
| | organismal development | organismal development | hematological system development and function |
| | cellular assembly and organization | nervous system development and function | lymphoid tissue structure and development |
| | tissue development | tissue development | cell-to-cell signaling and interaction |
| LUNG | amino acid metabolism | cellular function and maintenance | embryonic development |
| | cell death and survival | cellular movement | hair and skin development and function |
| | molecular transport | cell-to-cell signaling and interaction | organ development |
| | small molecule biochemistry | cell cycle | organismal development |
| | carbohydrate metabolism | cellular assembly and organization | tissue development |

**Table S4.** The summarization of the number of patients across subtypes identified by the clustering of ECv matrix and RNA-seq data.

| | | | RNA-seq | | | |
|---|---|---|---|---|---|---|
| | | | subtype 1 | subtype 2 | subtype 3 | all |
| ECv | subtype 1 | LUAD | 4 | 0 | 0 | 4 |
| | | LUSC | 224 | 0 | 0 | 224 |
| | subtype 2 | LUAD | 0 | 51 | 57 | 108 |
| | | LUSC | 0 | 10 | 3 | 13 |
| | subtype 3 | LUAD | 1 | 48 | 266 | 315 |
| | | LUSC | 3 | 7 | 18 | 28 |
| | all | | LUAD 5<br>LUSC 227 | LUAD 99<br>LUSC 17 | LUAD 323<br>LUSC 21 | LUAD 427<br>LUSC 265 |

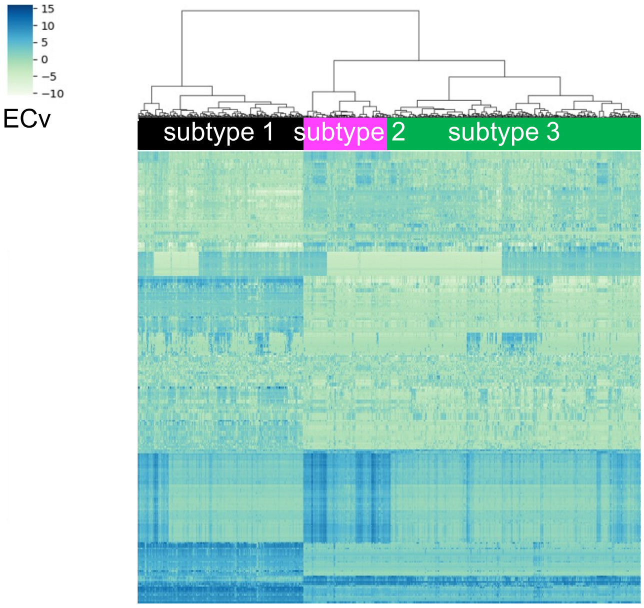


**Figure S1.** Heatmap showing hierarchical clustering for the ECv matrix in the LUNG dataset.

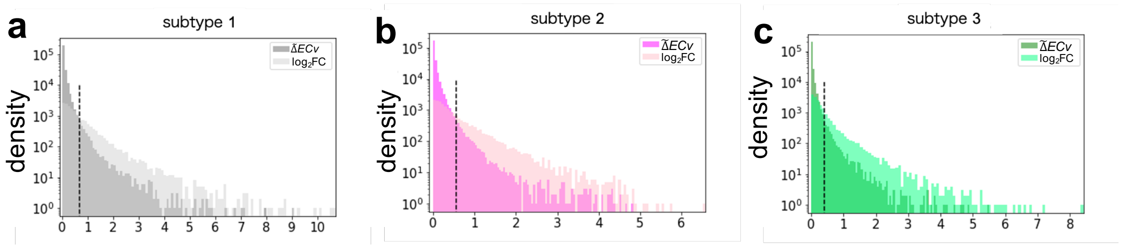


**Figure S2.** The distribution of $\tilde{\Delta}ECv$ of edges and $\log_2$ fold change (FC) in genes in the LUNG datasets. Dashed line represents the $\tilde{\Delta}ECv$ of the top 1.0% of total edges in every subtype.

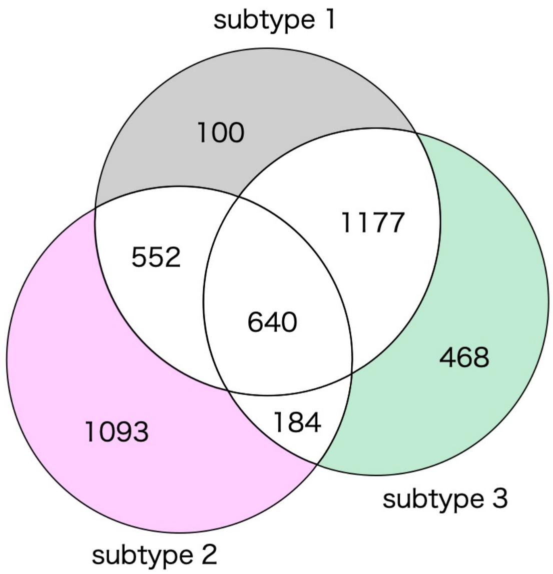


**Figure S3.** The Venn diagram represents the number of edges in the LUNG dataset. Colored areas in the Venn diagram represent subtype-specific edges in each subtype.

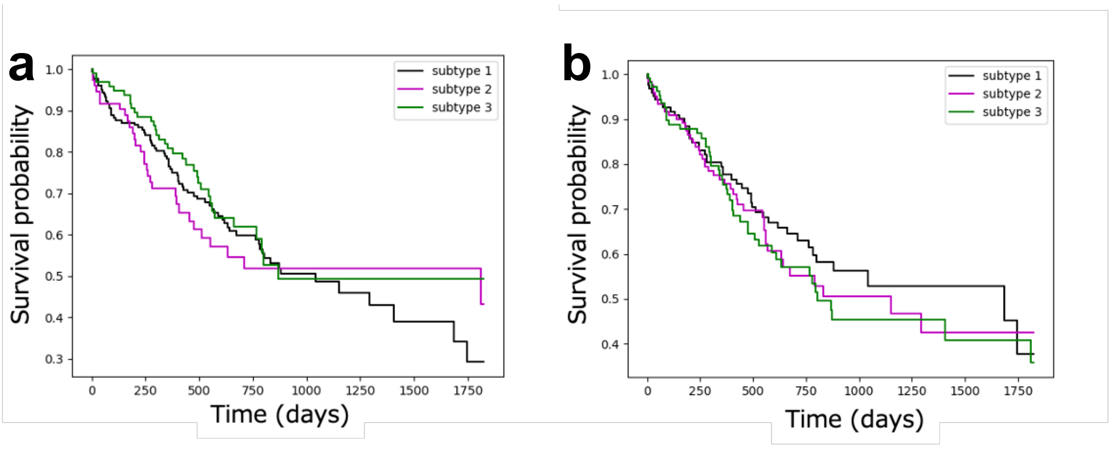


**Figure S4.** (a) Kaplan-Meier survival probability curves of patients identified by the iNMF method using gene expression data alone. The log rank test $\rho$-value between two subtypes; 0.93 (subtype 1 vs 2) > 0.05, 0.45 (subtype 2 vs 3) > 0.05, and 0.31 (subtype 2 vs 3) > 0.05. (b) Kaplan-Meier survival probability curves of patients identified by the iNMF method using multi-omics data. The log rank test $\rho$-value between two subtypes; 0.53 (subtype 1 vs 2) > 0.05, 0.34 (subtype 2 vs 3) > 0.05, and 0.78 (subtype 2 vs 3) > 0.05.

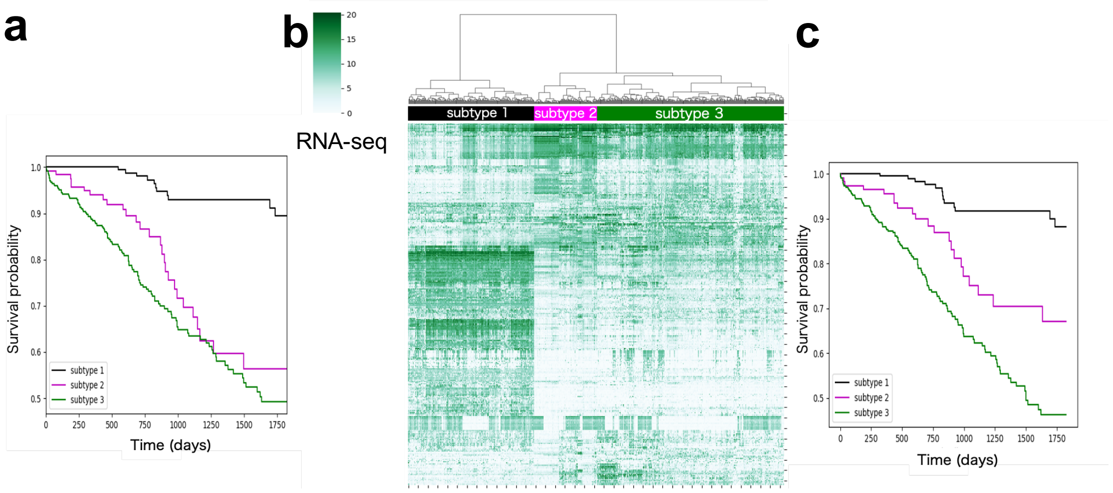


**Figure S5.** Analysis in the LUNG dataset. (a) Kaplan-Meier survival probability curves of patients for the identified ECv-based subtypes. The log rank test $\rho$-value between two subtypes; 3.1e-08 (subtype 1 vs 2) < 0.05, 9.6e-15 (subtype 1 vs 3) < 0.05, and 0.099 (subtype 2 vs 3) > 0.05. (b) Heatmap of the RNA-seq value matrix. (c) Kaplan-Meier survival probability curves of patients for the identified RNA-seq based subtypes. The log rank test $\rho$-value between two subtypes; 9.8e-05 (subtype 1 vs 2) < 0.05, 5.5e-15 (subtype 1 vs 3) < 0.05, and 3.6e-03 (subtype 2 vs 3) < 0.05.

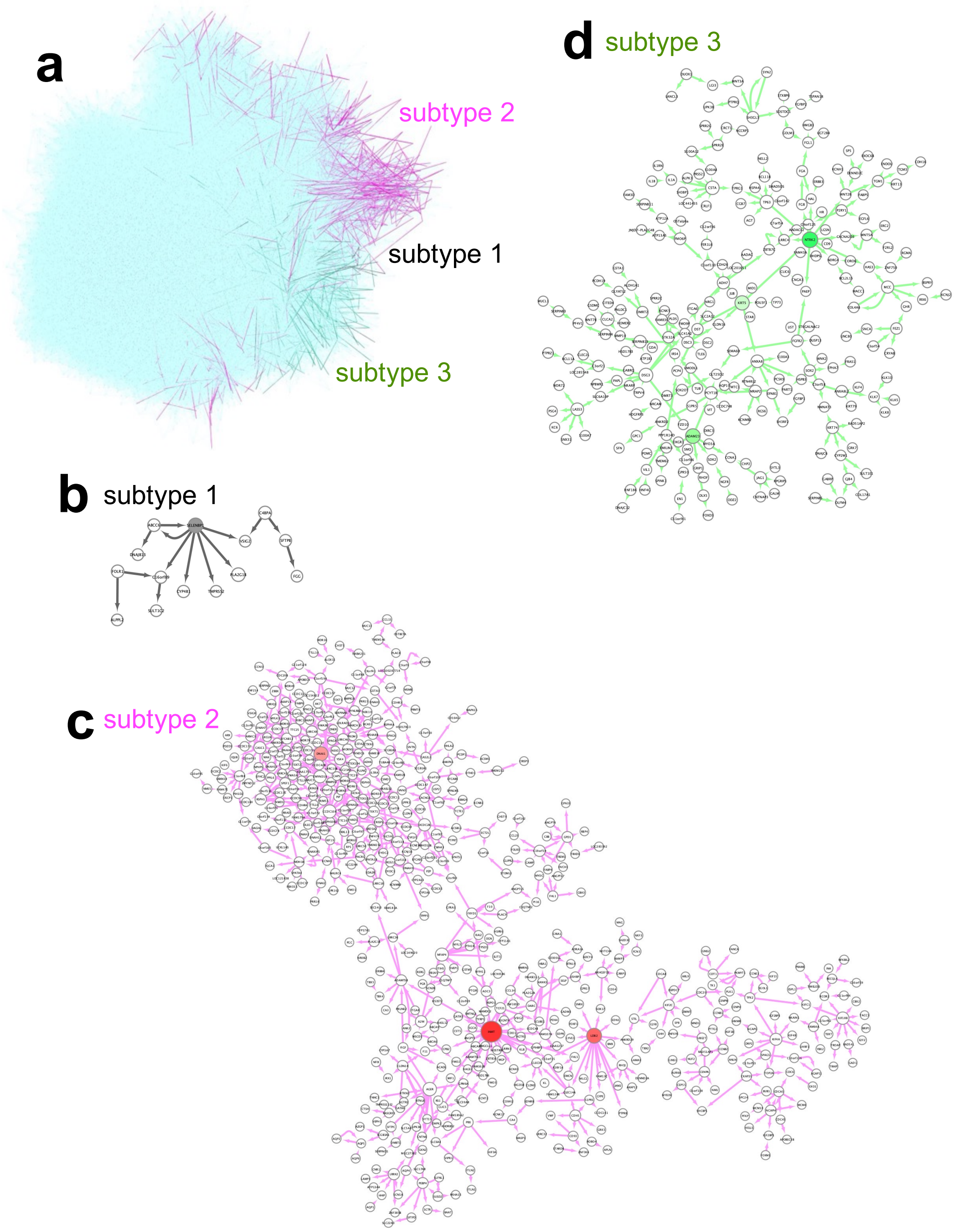


**Figure S6.** Visualization of subtype-specific subnetworks in the LUNG datasets.

(a) Subnetworks of subtype-specific edges were highlighted with the basal network (blue). (b-d) The biggest component in the subnetwork of subtype-specific edges in each subtype. Edges and nodes were colored by each subtype; subtype 1 (gray), subtype 2 (magenta), and subtype 3 (green). Colored nodes were hub nodes in each subtype and the color gradient represents the outdegree of hubs.
